# A Human Colon-based Microphysiological System with Mechanical Microenvironment Control for 3D Imaging

**DOI:** 10.1101/2025.04.10.648106

**Authors:** Duvan Rojas-Garcia, Dimitri Hamel, Julie Foncy, Dior Sambe, Sandra Pefaure, Laurent Malaquin, Audrey Ferrand

## Abstract

In vitro artificial colonic micro-devices that better replicate complex *in vivo* systems are essential tools for advancing our understanding of human gut (patho)physiology. Micro-physiological systems (MPS) offer controlled environments, enabling precise manipulation of tissue topography, rigidity and nutrient flow. We present the EnView system configured as a human colon-based MPS that faithfully replicates the 3D topography and matrix stiffness of the human colonic environment using an interpenetrating network of polyacrylamide and collagen I hydrogels, and enabling the culture of human colonic epithelium for weeks. The EnView system integrates a microfluidic chamber with active control of apical/luminal and basal/stromal compartments, allowing for in situ imaging and monitoring. Human colonic epithelial Caco-2 cells cultured up to 21 days in this system follow the crypt topography and formed a polarized epithelial monolayer.

This innovative MPS recapitulates in vitro a human colon epithelium with its 3D matrix topology and stiffness control, integrates a microfluidic chamber allowing active control of the apical/luminal and basal/stromal compartments by accurate injection, as well as in situ imaging.

**Teaser:** Human gut-on-chip reproducing tissue mechanical properties with active microfluidic control for 3D tissue characterization.

## Introduction

The colon corresponds to the distal part of the digestive tract. It plays a crucial role in the absorption of water, electrolytes, and nutrients, but also in the elimination of solid waste from the body, in bacterial fermentation, and in the synthesis of vitamins [1]. Absorption is ensured by the intestinal epithelium, consisting of a cell monolayer organized into two compartments: invaginations, called crypts, at the bottom of which are found intestinal stem cells (ISCs), and plateaus where differentiated cells (enterocytes, goblet cells, enteroendocrine cells, etc.) reside. Intestinal stem cells (ISCs) maintain the tissue by entirely renewing the epithelial cell monolayer within a week. Under physiological conditions, the intestinal niche controls crypt homeostasis and integrity [2]. This niche comprises the extracellular matrix (ECM), which regulates cell fate through matrix-cell interactions and its mechanical properties [3]. Indeed, cells perceive the mechanical properties of the matrix microenvironment (topography, stiffness, elasticity) and thus adapt their phenotype and behavior [4]. Alteration of the niche, particularly that of the mechanical properties of the extracellular matrix (ECM), can modify the regulation of ISCs and lead to the development of diseases such as inflammatory bowel disease (IBD) or cancer. In humans, unlike the small intestine, the colon is the frequent site of most pathologies such as Crohn’s disease, ulcerative colitis and cancer. Accurate modeling of the intestinal environment is crucial to better understand the initiation and development of intestinal diseases and to consider effective treatments.

For decades, intestinal (patho)physiology has been studied primarily using mouse models. The use of chemical agents (e.g., azoxymethane, DSS, TNBS) on mice or transgenic models have been used to study the pathogenesis of IBD or cancer [5, 6]. However, none reproduces the complexity of human pathologies, particularly regarding drug response and toxicity [7, 8]. For example, transgenic mouse models of cancer display disease mainly in the small intestine, whereas human intestinal cancer almost exclusively affects the colon (a.k.a. colorectal cancer). Researchers are therefore striving to develop more reliable in vitro cell culture models that can complement animal studies and better recapitulate human (patho)physiology. Although the intestine has been widely studied using in vitro models, these simplified systems often lack physiological relevance. Two-dimensional (2D) cell culture, used in the majority of in vitro studies, is inherently limited, as it is unable to reproduce the three-dimensional (3D) structure and interactions present in the native tissues.

In recent years, significant progress has been made in developing 3D models and dynamic culture systems derived from patient tissues or cells. These advanced models have become essential tools for investigating intestinal physiology, disease mechanisms, and drug responses. Notably, systems that mimic the structural complexity and multicellular composition of the intestinal barrier have gained increasing attention and widespread use. [9]. Recent advancements in bioengineering have led to the development of new in vitro culture models aiming to more faithfully reproduce tissue physio(patho)logy by controlling multiple parameters of the cellular microenvironment. These devices are named microphysiological systems (MPS). The International Consortium for Innovation and Quality in Pharmaceutical Development (IQ) defines them as “culture systems that go beyond traditional 2D culture including several of the following design aspects: a multicellular environment within a biopolymer or tissue-derived matrix, a 3D structure, the inclusion of mechanical cues such as stretch or infusion, the incorporation of primary cells or derived from stem cells, and/or inclusion of immune system components” [10]. These MPS can be advantageously coupled to a microfluidic system, which in the case of intestinal systems, makes possible to access and control mass transport the ‘luminal’ and/or ‘stromal’ compartments. The fine control of luminal flow makes it possible to mimic in vitro the passage of material in the lumen of the digestive tract, which makes this type of model very relevant in the study of the digestive tract.

In recent years, various MPS configurations have been developed targeting different aspects of the intestinal epithelium. For instance, MPS coupled to fluidic flow, thus inducing shear stress on cells, use luminal and basal chambers separated by a porous membrane, allowing 2D culture of epithelial cells and control of the apico-basal flow [11]. Additionally, two hollow lateral chambers have been integrated, allowing control of air pressure and thus stretching or compression of the central chamber membrane [12, 13]. Alternatively, some models use biocompatible hydrogels - either synthetic or natural - whose porosity and capacity allow small molecules to diffuse freely, thus promoting the formation of growth factors, oxygen and cellular waste gradients, essential for tissue homeostasis. For example, Type I Collagen (Coll I) has been used instead of a membrane to create a hydrogel interface separating fluidic channels [14].

Other approaches have focused on replicating 3D intestinal topography (crypt and/or villi) and an accessible lumen/epithelium interface. For example, Coll I hydrogel scaffolds were used to culture murine intestinal organoids and fibroblasts in the gel [15]. Similarly, other studies have presented intestinal epithelial models using murine organoids on a Coll I mixed with Matrigel by molding respective tissue geometries (i.e. stomach, small intestine, caecum and colon), and have highlighted stem cell compartmentalization and transcriptional resemblance to epithelia [16]. Other studies have used photopolymerizable hydrogels (PEG-DA, gelatin methacryloyl (GelMa) or silk fibroin-gelatin (SF-G) based hydrogels), in combination with high-resolution, laser-based 3D printing, enabling the growth and organization of intestinal cell lines within structured environments [17, 18]. In all cases, the groups demonstrated the impact of topography on the establishment of a uniform epithelium, cell polarization and migration. A similar approach used a PDMS replication technique, where the murine colon crypt topography was patterned into a Coll I hydrogel layer positioned on the membrane of a transwell-type insert. This configuration allowed the establishment of a biochemical gradient along the crypt axis by placing different media compartments (i.e. basal and luminal) [19], as well as an oxygen gradient to study co-cultures of the human gut microbiome, and more recently, fibroblast incorporation [20, 21].

Some models have combined both active fluidic flow and three-dimensional support to control the dynamics and spatial distribution of mass transport in the tissue. For example, Shim *et al*. cultured Caco-2 cells on a PDMS chip containing a collagen scaffold with 3D villi supported on a porous membrane, showing an increased activity of one of the drug metabolism enzymes, cytochrome P450 3A4, when the cells were cultured in 3D under flow compared to 2D cultures [22]. As an alternative, another colon MPS has been developed, presenting a channel with crypt-like structures that reproduces a ‘lumen’, formed by laser ablation on a hybrid hydrogel (mixture of Coll I and Matrigel). This system was coupled to a fluidic system allowing active control of the luminal compartment above the colonic epithelium derived from human organoids [23, 24]. Moreover, on the basal side of the epithelial linen, the culture medium was supplied by diffusion from a reservoir. Similarly, Vera *et al*. presented an intestinal MPS model featuring a channel with villus-like structures by 3D bioprinting using gelatin-methacrylate (GelMA) and polyethylene glycol diacrylate (PEGDA) [25]. This MPS was validated using Caco-2 cells and allowed measurements of luminal and basal flow, as well as trans-epithelial electrical resistance (TEER).

However, to our knowledge, only a few models have been developed not only to recapitulate 3D tissue architecture and active flow control of both luminal and basal compartments, but also account for the stiffness of the matrix material. Yet, matrix stiffness is a critical parameter, as human colon tissue exhibits distinct mechanical properties under physiological conditions and pathological settings, with stiffness ranging from 1–3 kPa [26, 27] to 8–28 kPa in IBD (Crohn’s disease and ulcerative colitis) [28] and up to 68 kPa in colorectal cancer [27]. This alteration in matrix stiffness is partly due to increased collagen cross-linking and decreased matrix degradation [29]. Thus, to mimic accurately the complex architecture and functional cues of colon tissue, an effective MPS should integrate key features such as precise 3D replication of human colon crypt topography, matrix stiffness control to reflect physiological or pathological conditions, and a biomimetic extracellular matrix that supports cell adhesion and differentiation. In addition, the MPS must allow for the establishment of apical-basal polarity, fluidic flow to simulate nutrient and waste transport, while establishing biochemical and oxygen gradients to sustain the tissue microenvironment.

Here, we present the development and characterization of an alternative human colon microphysiological system (MPS) designed as an imaging-compatible chamber integrated with a microfluidic platform (Figure 1). This system was engineered to precisely control colon tissue topography, tune the stiffness of the supporting scaffold, manage apical/luminal and basal/stromal compartments through accurate active fluidic injection and sample recovery, and enable high-resolution in situ imaging.

**Figure 1.**
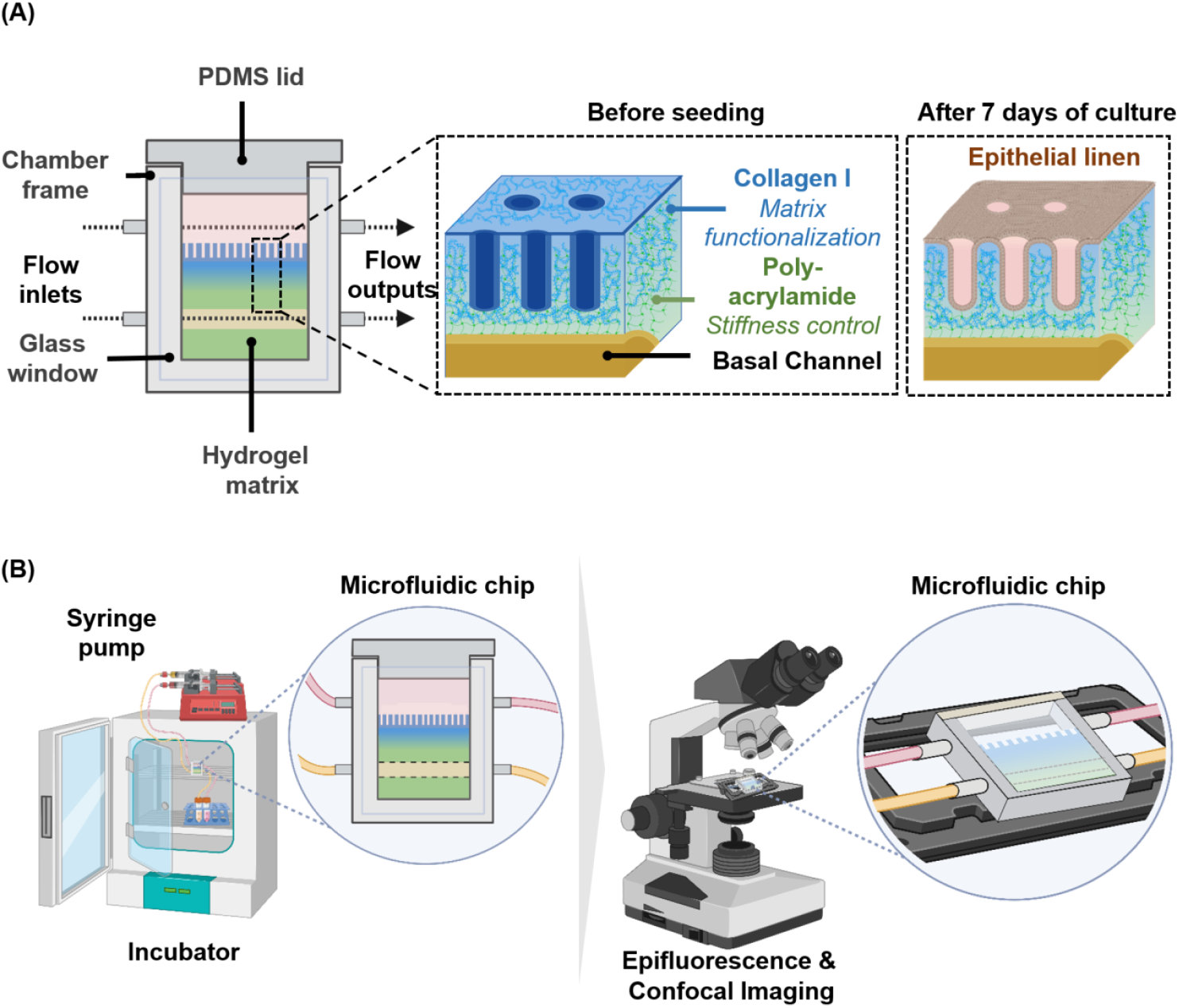
Schematic representation of the EnView colon microphysiological system (MPS): (A) Human crypt-based structures were molded in an interpenetrating hydrogel (IPN) of polyacrylamide and type I collagen (Coll I) using a 3D-printed mold (high-resolution DWS29J+ printer). The hydrogel combines the stiffness control and reproducibility of IPN hydrogels, and the cell attachment and growth capabilities of collagen I. The device is enclosed in a closed stainless-steel chamber composed of two distinct compartments, an upper ‘luminal’ compartment and a lower stromal-like compartment. (B) Schematic representation of the experimental setup. The EnView microphysiological system is placed inside an incubator (37 °C, 5% CO2). The luminal and stromal environments can be actively controlled by a microfluidic pump, allowing perfusion of two different cell culture media. The platform easily enables in situ tissue imaging, as the EnView microfluidic chamber functions as a microscopy chamber.

The EnView system introduces an original chip architecture that integrates a microfluidic frame with transparent lateral windows, optimized for three-dimensional optical imaging. Its fabrication follows a three-step process (Figure 2): first, assembling the chamber by mounting two glass slides onto a rigid frame that incorporates the microfluidic channels; second, molding biomimetic crypt-like structures into a hydrogel with defined mechanical properties using a 3D-printed mold; and third, integrating luminal and basal compartments to establish a fully functional microfluidic system capable of dynamic and spatial control of molecular transport.

**Figure 2.**
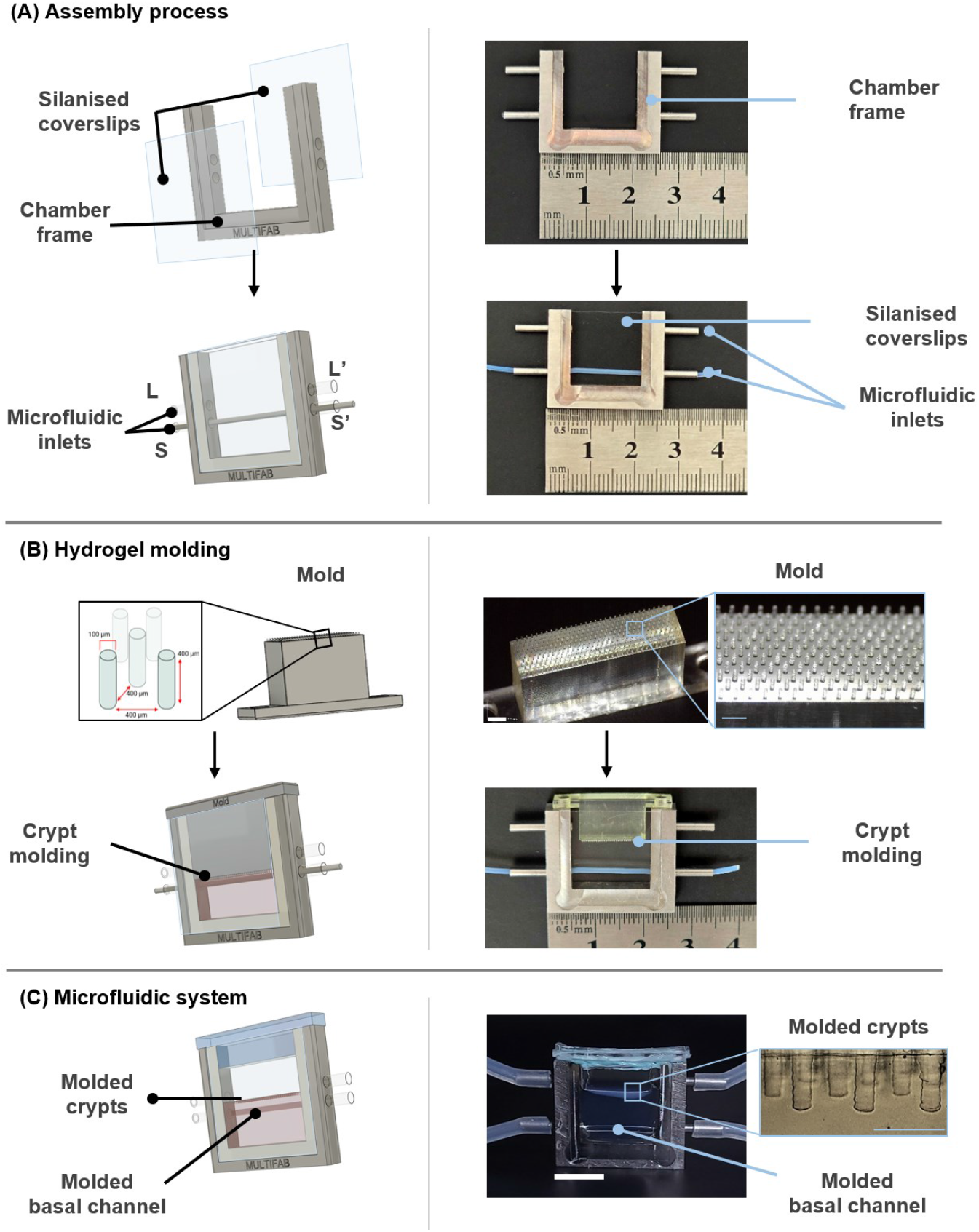
Fabrication workflow of the EnView colon microphysiological system (MPS): (A) A stainless steel U-frame is assembled with two silanized glass side coverslips and tube segments (luminal: L and L’, stromal: S and S’). A microfluidic tube is inserted to mold the stromal/basal channel during hydrogel polymerization. Scale bar: 1 cm. (B) A 3D-printed mold coated with a layer of cross-linked collagen I (5 mg/ml), comprising an array of eleven 400 µm-high, 100 µm-diameter pillars arranged in a hexagonal lattice (400 µm period, 11 rows), is inserted into the chamber containing the PA hydrogel preparation. Scale bar of the mold: 1000 µm. (C) After polymerization, the microfluidic tube and mold are removed, and the device is sealed with a PDMS lid. Scale bar of the frame (white): 1 cm, and scale bar of the hydrogel (light blue): 500 µm.

To recreate the complex topography of the human colon epithelium, we employed high-resolution molding of the hydrogel using 3D-printed templates. The hydrogel is based on an interpenetrating polymer network (IPN) composed of polyacrylamide and type I collagen (Coll I). IPNs consist of two or more interwoven, functionally distinct polymer networks that are physically entangled without covalent bonds. This structure enables independent tuning of mechanical and biochemical properties, offering a physiologically relevant microenvironment for cell culture. In our system, the IPN hydrogel provides adjustable stiffness and enhanced structural integrity, while also supporting cell adhesion and proliferation. The molding process is performed directly within the EnView microfluidic device, which features multiple inlet and outlet channels to allow continuous perfusion during cell culture. Importantly, the EnView device includes two lateral glass windows, offering unique access for real-time, high-resolution imaging of tissue formation and development throughout the culture period.

In this study, we first investigated the resolution of the molding process associated with PA varying compositions. We used two preparations targeting physiological and pathological stiffness of 3.8±1.8 kPa and 26.2±8.4 kPa, respectively, which we will refer to as soft and hard materials. We then characterized the molded crypts obtained in the hydrogels, verifying the reproducibility of the fabrication, the formation of Coll I fiber network, and molecular diffusion. Finally, we established Caco-2 cell culture conditions allowing the establishment of a monolayer of human colorectal epithelium extending from the bottom of the crypts to the planar interface of the scaffold. The epithelial constructs were polarized in 3D at 14 days of culture.

## Results

### Replication of the human colon crypt topologies into the IPN hydrogels

A first step was to determine the overall stiffness of the hydrogels that we wish to use within the device (Supplementary Figure 1). A liquid formulation of Collagen I was prepared at 5mg/ml and reticulated at 37°C. Two mixes of polyacrylamide were prepared changing acrylamide:bis-acrylamide ratio, mix 1 corresponding to ratios of (1: 1.12) and mix 2 to (1: 0.45), and then charged onto the Coll I hydrogels. Compression measurements were performed on the two polyacrylamide (PA) and Collagen I IPN hydrogels of different stiffness confirming that we can produce IPN hydrogels composed of PA and Coll I with stiffness values of 3.8±1.8 kPa and 26.2±8.4 kPa, respectively reproducing the physiological or pathological (inflammatory) stiffness of the human colon.

The next experiments were devoted to the molding of human colon-like crypt structures in the IPN hydrogels. A liquid formulation of Coll I at 5 mg/mL was deposited on the 3D printed mold. After reticulation at 37°C during 30 minutes, the mold was inserted into the EnView chamber containing a liquid PA formulation. Dimensional analysis (Figure 3) of the height and diameter of the crypt structures showed similar mean diameter values of 82.1 ± 11.9 µm and 75.3 ± 9.6 µm for the soft and hard materials respectively (n=3). The mean height of the structures was estimated to 303.6 ± 7.4 µm and 345.1 ± 4.8 µm, respectively. These values are of the order of the dimensions of the crypts of the human colon: 70-100 µm in diameter and 320-450 µm in height [31] and are in good agreement with the expected mold dimensions (Supplementary figure 2), although slightly smaller, 98.2 ± 5.0 µm in diameter and 417.7 ± 9.8 µm. We observed a difference of approximately 16% and 23% in diameter and 27% and 17% in height for soft and hard hydrogels respectively.

**Figure 3.**
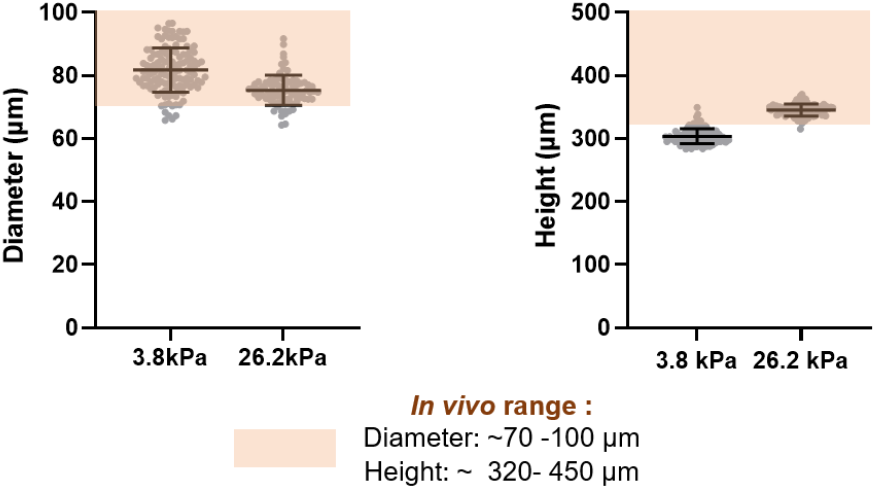
Characterization of the molded crypt structures: Measurements of crypt structures within 3.8 kPa and 26.2 kPa IPNs from bright-field images after hydrogel casting on MPS (n = 3). Orange rectangles represent the physiological dimensions of crypts in vivo: 70–100 µm in diameter and 320–450 µm in height (42).

The differences between the mold dimensions and the molded crypts could result from a swelling mechanism that may be associated with the PA reticulation process during molding [32]. The higher acrylamide concentration in hard hydrogels resulted in the formation of a highly cross-linked rigid structure where the gel network has restricted mobility and does not absorb as much water as in soft hydrogels [33]. Consistent with this first observation, the swelling mechanism is also likely to induce a slight increase in the observed gap between pillars, estimated at 427 and 420 µm for soft and hard materials respectively. Importantly, the crypt dimensions are reproducible throughout fabrication batches (n=3) for both stiffness.

### Characterization of Coll I/PA IPN hydrogels

We further investigated the formation of Coll I fibers in the hydrogel, which enables cell adhesion, as PA alone cannot promote cell adhesion to its surface. Using second harmonic generation (SHG) imaging, we observed that the mold pillar structure (Supplementary figure 2) was successfully replicated in the IPN hydrogel (Figure 4A). Furthermore, in vitro reticulation of Coll I enabled the formation of fibrillar networks (for another illustration of the fibrillar network, see supplementary figure 3) [34].

**Figure 4.**
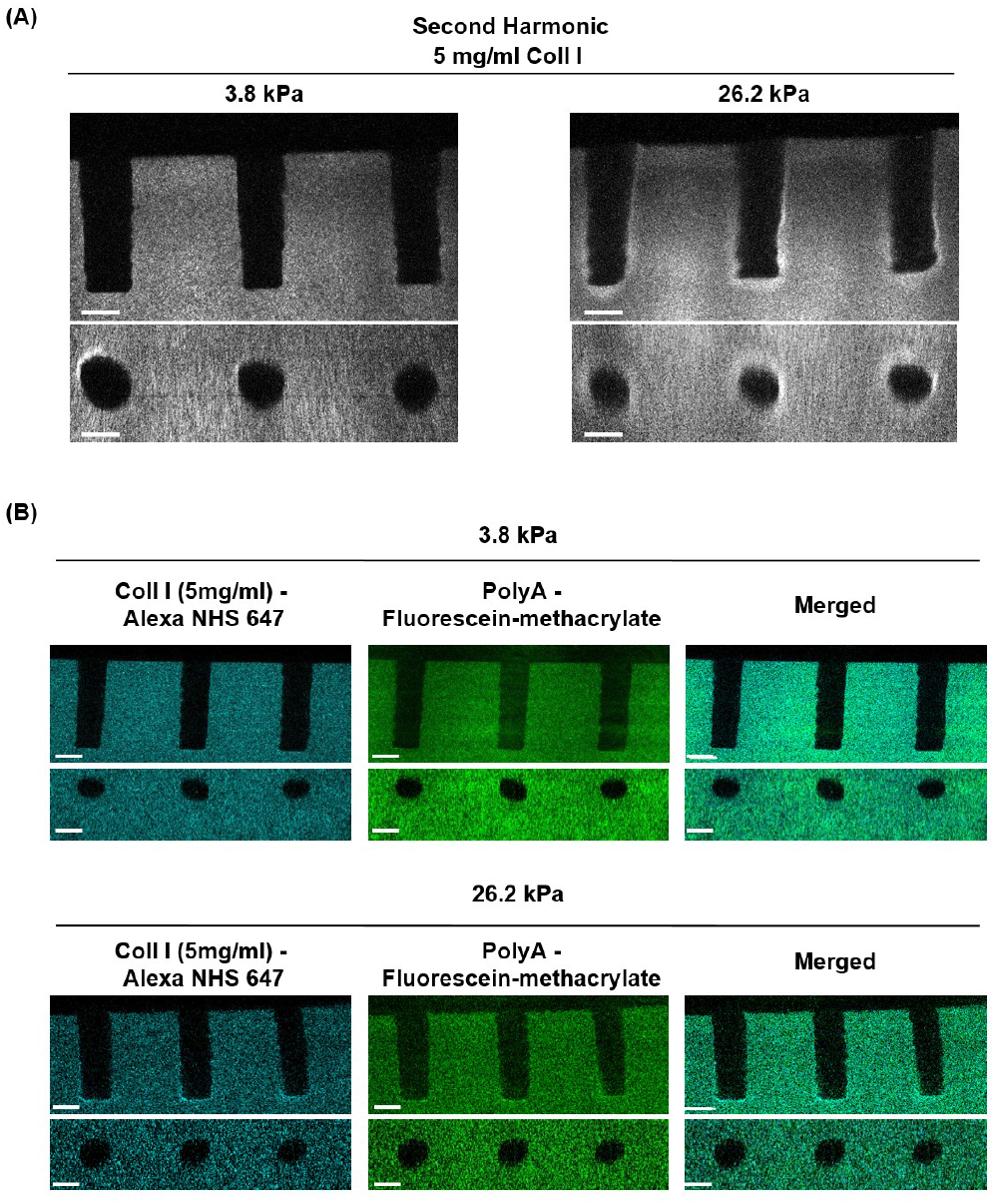
Interpenetrating hydrogel (Collagen I and Polyacrylamide) characterization: (A) Second harmonic imaging of collagen I network. Scale bars: 100µm. (B) Confocal imaging of stained interpenetrating hydrogels: Polyacrylamide (fluorescein-methacrylate, green) and Collagen I (AlexaFluor 647 NHS, cyan). Homogenous interpenetration of polyacrylamide and collagen I was obtained for both stiffness. Scale bar: 100µm.

To assess the spatial distribution of Collagen I (Coll I) and polyacrylamide (PA) within the hydrogel, we selectively labeled each component prior to crosslinking: fluorescein dimethacrylate for PA and AlexaFluor 647 NHS (N-hydroxysuccinimide) Ester for Coll I. Fluorescein dimethacrylate binds to polyacrylamide (PA) through covalent bonds with the PA network during polymerization, creating a stable, fluorescently labeled matrix. AlexaFluor 647 NHS Ester is a fluorescent dye that binds to primary amines on proteins (such as collagen) through an NHS Ester bond. Together, these dyes allow for the visualization and spatial mapping of each hydrogel component.

Intensity profiles (Figure 4B) show a similar spatial distribution of both hydrogels (PA and Coll I), confirming the existence of an IPN hydrogel of both materials in the area where human crypt-like structures are molded.

### Molecular diffusion characterization trough hydrogels

To assess the perfusion of molecules through the stromal channel, a crucial parameter for basal feeding of cell cultures in culture medium, we investigated the diffusion capacity of molecules of different molecular weights through hydrogels in MPS. We injected fluorescein isothiocyanate (FITC)-labeled dextran solutions of different molecular weights (3 kDa, 70 kDa, and 150 kDa) into the basal channel at a flow rate of 100 µL/min, ensuring a complete renewal of FITC-dextran in the basal channel every minute. The time evolution of the molecular diffusion profile of the fluorescent solutions was characterized by confocal microscopy. We considered the normalized gray values profiles of fluorescence intensity as representative of the concentration profiles. Figure 5A shows, as expected, faster diffusion in soft (3.8 kPa) hydrogel compared to hard (26.2 kPa) formulation. This observation was consistent across all molecular weights tested. As polymer concentration increases, the resulting PA network becomes denser, restricting molecular mobility and reducing mass transport by diffusion. Similarly, when increasing the molecular weight of dextran molecules, we observed a significant decrease of diffusion rates for both stiffness. When increasing from 3 to 150kDa, the diffusion coefficient values drop from 94.4 ± 28.6 µm^2^/s to 1.3 ± 0.6 µm^2^/s and from 44.6 ± 10.1 µm^2^/s to 0.2 ± 0.2 µm^2^/s, for soft and hard hydrogels (n=3), respectively. These results are explained by an increase in the hydrodynamic diameter of molecules at higher molecular weights, which, as established by the Stokes-Einstein equation, is inversely proportional to the diffusion coefficient [35].

**Figure 5.**
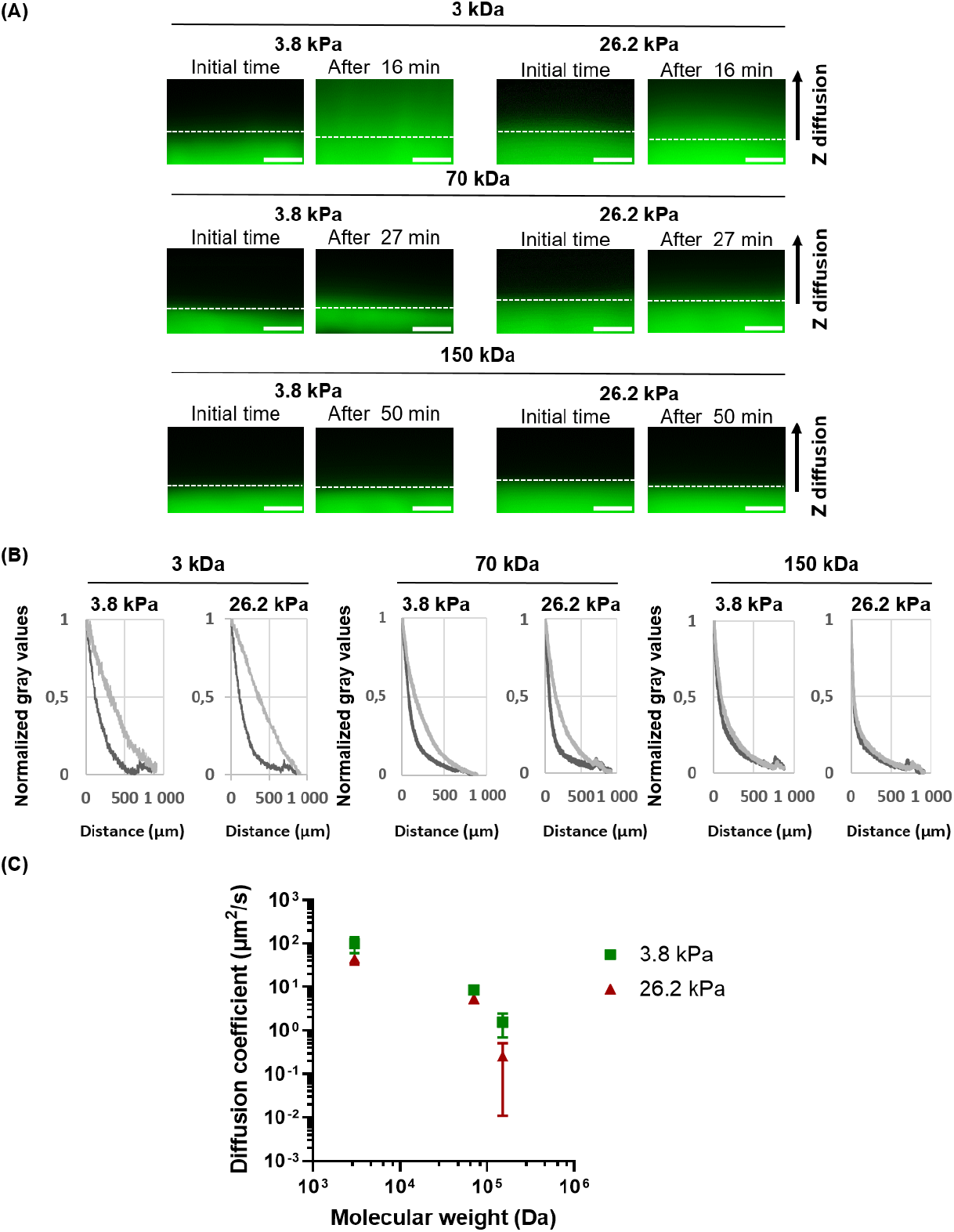
Diffusion characterisation through an IPN hydrogel at two different stiffness (3.8kPa and 26.2kPa). (A) FITC-dextran solutions of different molecular weights were injected in the stromal/basal channel (upper limit represented by white dotted line) of the MPS at a 100µL/min flow rate. Acquisition time was defined in order to obtain similar distances of diffusion front displacement. Pictures were taken every 10 seconds. Scale bars: 500µm. (B) Normalized gray values were plotted against distance in order to compare diffusion profiles of the two stiffness conditions. (C) Diffusion coefficients plotted against molecular FITC-Dextran molecular weights. Mean and SD values are presented (n=3).

Interestingly a threshold value was found for both hydrogel formulations at 150kDa. For FITC-dextran molecules with higher molecular weights (150kDa), we observed a very slight evolution of the concentration profile over time, suggesting very low molecular transport in the hydrogels (Figure 5B), thus validating previous observations. Our findings are in line with previous observations of FITC-Dextran diffusions reported in PBS for similar molecular weights (4.4 and 172.5 kDa), 127.7 µm2/s and 36.5 µm2/s respectively [36]. Overall, these results suggest that molecules such as cell growth factor or cytokines, whose molecular weights are generally between 6 and 70kDa [37], would be able to diffuse from the basal channel though the PA hydrogel.

### Colorectal culture into the human colon MPS

A crucial aspect of this study was to establish cell culture conditions that would support the formation of a continuous epithelial lining from the crypt bases to the surface of the plateaus. To achieve this, we plated Caco-2 cells onto devices containing either a soft or a hard IPN hydrogel. Bright-field images were taken at different times during culture. Caco-2 cultures were stopped at day 7, 14 and 21 in order to perform characterizations by immunofluorescence staining to observe nuclei (DAPI, cyan) and actin network (Phalloidin, magenta). Immediately after plating, cells were present at the bottom of the crypts and on the plateau zones (data not shown). Epithelial cells successfully adhered and colonized the crypts and the plateaus of the scaffold by day 7, forming a continuous epithelial monolayer onto both soft and hard hydrogels (Figures 6 and 7). In some instances, when multiple cells were present in the upper region of the crypt, they extended across the opening rather than migrating along the crypt wall (data not shown). This behavior has also been observed for Caco-2 cells cultured on crypt-patterned PDMS substrates [38].

**Figure 6.**
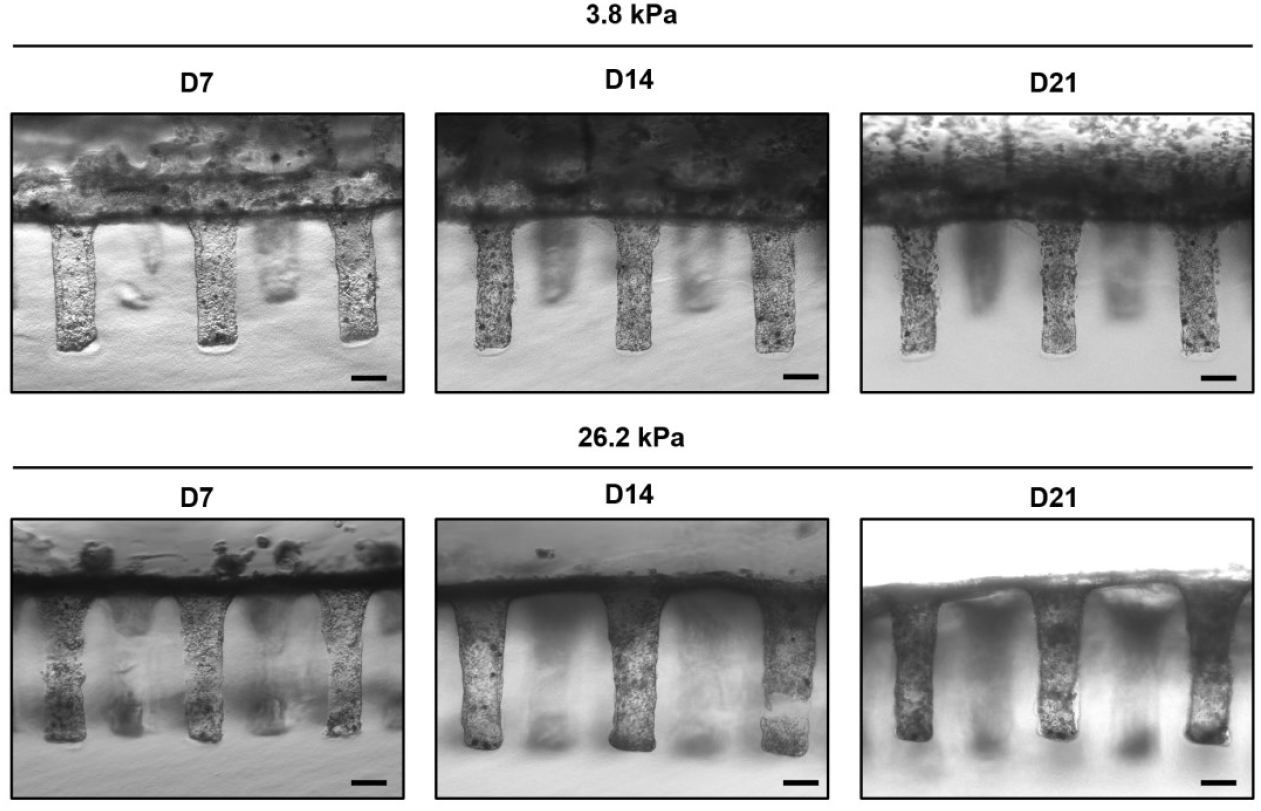
Caco-2 cell line culture on the colon MPS during 21 days: Follow-up bright-field images of Caco-2 cell line culture on 3.8kPa and 26.2 kPa matrices. Culture was kept for 21 days and imaged at day 7, 14 and 21. Scale bars: 100 µm.

**Figure 7.**
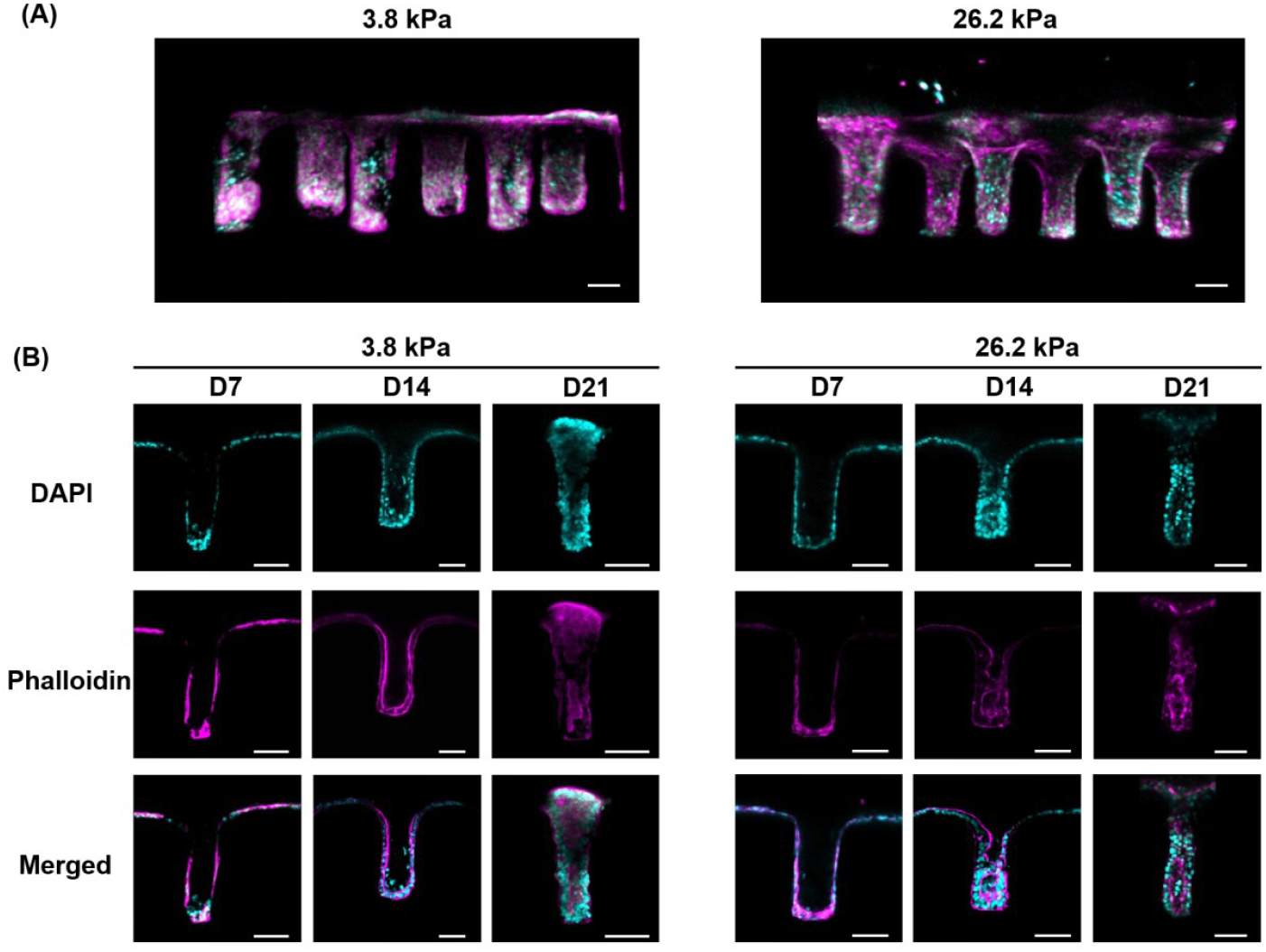
Caco-2 cell line culture on the colon MPS Immunostaining characterization: Cells were cultured on 3.8 kPa and 26.2 kPa IPN hydrogels. Cells were fixed and stained for DNA (DAPI, cyan) and F-actin (Phalloidin, magenta). Images were acquired by confocal microscopy. (A) Large 3D view (10X) of culture cultures at day 14. (B) Z stacks for crypt comparison between cultures stopped after 7 days, 14 days, and 21 days respectively. Scale bars: 100 µm.

After 14 days of culture, we were able to visualize a thickening of the epithelial wall for cultures established on hard IPN hydrogels while this event was observed later (day 21) for cultures on soft hydrogel, suggesting that a harder scaffold stiffness favors the acquisition of polarization. Moreover, at day 21, we could observed crypt hyperplasia on hard hydrogels, cells forming a multilayer in certain parts of the crypt, as seen in Figure 7.

## Discussion

We demonstrated a simple and efficient method for the fabrication of a human colon MPS system combining topology, the matrix scaffold stiffness control, the creation of a lumen and basal compartments, and a microfluidic circuit to control mass transport in the model. A first proof of concept was validated using a hydrogel composed of a polyacrylamide support layer under an IPN of polyacrylamide and collagen I. We first hypothesized that the addition of cross-linked Coll I to the chamber containing liquid polyacrylamide promotes the creation of an IPN (Figure 1). This was confirmed by image analysis of the material distribution that shows the overlap of the two materials, especially in the crypts area. We also believe that the accuracy of replication improves with increasing material stiffness. It is known from the literature [39] that the stiffness of Coll I, in the concentration range of 1 to 5 mg/mL, exhibits Young’s modulus values, varying from 100 Pa to 1.4 kPa, respectively. We can reasonably estimate that the stiffness of the Coll I/PA IPN is thus slightly influenced by the properties of the collagen hydrogel. This observation converges with the evolution of the structure dimensions which suggests a better replication accuracy for the highest polyacrylamide concentrations. This trend was confirmed by the analysis of the structure diameter, which presents the best accuracy for the combination Coll I 5 mg/mL and hard PA. This phenomenon also corresponds to the evolution of the structure height. This method clearly opens the way towards the creation of heterogeneous matrices reproducing the tissue architecture and gradients of factors and stiffness found *in vivo*. Indeed, the design of this MPS could make possible the integration and structuration of a large variety of both synthetic and natural hydrogel materials (*e*.*g*. collagen, gelatin, laminin). The system could also be compatible with current bioprinting systems to create heterogeneous models of ECM.

Next, we experimentally determined diffusion coefficients in 3.8 kPa and 26.2 kPa hydrogels using a range of fluorescein isothiocyanate (FITC)-labeled dextran solutions (average molecular weight 3, 70, 150 kDa). Molecules relevant to cell culture up to a molecular weight of 70 kDa, such as cell growth factors or cytokines for example, were reported to be able to diffuse from the basal channel through the PA hydrogel and reach the cells. However, based on our results, it is likely that species with molecular weight around and higher than 150kDa, would not be able to pass through the hydrogel by diffusion. We therefore conclude that immunostaining of the epithelium in our system must be performed via the luminal compartment, following tissue permeabilization when necessary. In this context, the microfluidic interface—enabling precise injection into and recovery from the system—offers the possibility of perfusing different media. This capability could be leveraged to establish growth factor gradients, thereby promoting the compartmentalization of primary cell cultures. The microfluidic set-up will also made possible the real time control and monitoring of the culture medium composition in both compartments via injection and enable the recovery of factors, markers, etc. This later aspect is particularly interesting for the follow-up of the cell secretome and metabolism. Getting access to the different MPS compartments could allow exploring interactions between the luminal content, which could include microbiota, nutrients, food additive or contaminants, and the colorectal epithelium.

Furthermore, we successfully demonstrated the possibility of establishing an epithelial lining on this hydrogel, while preserving the resolution and precision of the creation of the human colon architecture. Important criteria for us while designing this MPS was the possibility to study the epithelium by several complementary means. A specific attention was paid to the ease of observation and imaging of the content and, in particular, of the epithelial tissue. Thus, we integrated glass slides on either side of the MPS. The design of the mold was also optimized in such a way as to provide an optimal working distance to perform wide field and confocal imaging approaches, including high-resolution analysis. This allowed us to observe that our device allowed the culture of a colonic cell line (Caco-2) for 21 days, successfully reproducing the geometry and dimensions of human colonic crypts, while allowing imaging monitoring of culture establishment. Epithelial monolayers were polarized at 14 days, when it takes nearly 21 days for traditional 2D culture of these cells. This type of early polarization has also been previously reported in cultures subjected to physical constraints such as shear stress induced by medium perfusion [40]. We believe that an important feature of the proposed MPS relies in the possibility to create a large epithelial surface area of approximately 1 cm^2^ which is wider than most of the current systems described above. This notion is crucial as it would certainly allow to recover enough material to perform either transcriptomic or proteomic approaches.

In the future, this human colon MPS, combined with patient-derived colon organoids from individuals affected by conditions such as inflammatory bowel disease, metabolic disorders, or familial adenomatous polyposis, could significantly enhance the efficiency of screening platforms -for drugs, nutrients, dietary contaminants, and microbiota interactions. It could also serve as a powerful tool for testing vectorized drug delivery and for in vitro diagnostic or prognostic applications. Incorporating immune components and enabling co-culture with microbiota will further increase the physiological relevance of the model, better replicating native gut tissue functionality. Ultimately, this platform aligns with the 3R principles, supporting the development of alternative strategies to reduce reliance on animal experimentation.

## Materials and Methods

### Collagen I preparation

To prepare 1 mL of a neutralized collagen type 1 solution (at final concentration 5 mg/mL), 454.5 µL of collagen (Corning 354249; rat tail collagen type I, 11 mg/mL in 0.02 N acetic acid) were mixed with 10.4 µL of sodium hydroxide (NaOH; Sigma S2770; stock concentration of 1 mol/L, working concentration of 30 mmol/L), 20 µL of HEPES (Sigma 83264; stock concentration of 1 mol/L, working concentration of 20 mmol/L), 132.7 µL of sodium bicarbonate (NaHCO3; Sigma S5761; stock concentration of 7.5% wt/vol, working concentration of 0.58% wt/vol), 100 µL of 10X phosphate-buffered saline (PBS; Sigma D1408; working concentration of 1X), and 282.3 µL of deionized water on ice. This solution was then mixed by pipetting and briefly vortexing. To stain the collagen network, 0.3 µL AlexaFluor 647 NHS Ester (Life Technologies, A37573, stock concentration 0.1 mg/mL) were added to collagen I solution prior reticulation.

### Polyacrylamide preparation

To prepare 1 mL of PA solution, 250 µL for hard PA gel, or 100 µL for soft gel, of acrylamide (Sigma A3553; stock concentration of 40%, working concentration of 10% for hard PA gel or 4% for soft PA gel) were mixed with 112 µL of bis-acrylamide for both rigidity gels (N,N’Methylenebisacrylamide; Sigma M7279; stock concentration of 2%, working concentration of 0.22%), and 627 µL of deionized water for hard gel or 786 µL for soft gel. For PA hydrogel staining, 10 µL of fluorescein dimethacrylate (Sigma 570249; stock concentration of 1 mg/mL, working concentration of 10 µg/mL) were added. Reticulation reaction was started by extemporaneously adding the following catalyzers: 10 µL APS (Ammonium persulfate; Sigma 248614; stock concentration of 10%, working concentration of 0.1%) and 1 µL TEMED (Tetramethylethylenediamine; Sigma T7024; working concentration of 0.1%). The mix was immediately vortexed and seeded in the device chamber.

### Glass silanization

To create covalent bounds between polyacrylamide hydrogel and glass in our microdevice, we performed glass coverslip treatment according with a usual APTES/glutaraldehyde protocol [41]. Glass coverslips (20mm x 20mm) were first washed with acetone, deionized water and ethanol before dried. Then, coverslips were treated with oxygen plasma for 5 minutes (100% generator power and 0.5mbar pressure, Plasma Pico system, Diener GmbH). Plasma treated coverslips’ sides were incubated in pure APTES ((3-Aminopropyl) triethoxysilane; Sigma, 440410) solution during 15 minutes at room temperature and then washed three times with deionized water. Coverslips were then incubated for at least 30 minutes (maximum 1 hour) in glutaraldehyde 0.5% solution (Sigma, 340855 reconstituted in deionized water) protected from light and washed three times with 50 mL of deionized water before drying. Silanized glass coverslips could be stored on Parafilm® at 4°C in hermetic Petri dish during 3 weeks until use.

### IPN hydrogel preparation on silanized glass coverslips

To measure the stiffness of the IPN hydrogels, we developed an IPN hydrogel preparation directly onto silanized glass slides (supplementary figure 1). First, 30 µL of collagen type I (Coll I) solution was deposited onto a 3D-printed mold and polymerized at 37°C for 30 minutes. After this, 100 µL of either soft or hard PA solution was added over the polymerized collagen, and a silanized glass coverslip was placed on top of the mold. The setup was left to crosslink at room temperature for 30 minutes. Following polymerization, the mold was gently removed to reveal the crypt structures within the IPN hydrogel on the silanized glass coverslip. These structures were rinsed in phosphate-buffered saline (PBS) and stored at 4°C until use. The combination of Coll I and PA allowed for precise replication of crypt topography, promoting both structural integrity and cell adhesion. This approach facilitated the fabrication of a PA & Coll I IPN hydrogel that mimicked the 3D architecture of colonic crypts while supporting epithelial monolayer formation. It ensured reproducibility and provided a stable platform for subsequent compression measurements.

### Compression measurements

Compression tests were conducted to evaluate the stiffness of the hydrogel materials, a critical factor for mimicking physiological and pathological conditions. The setup involved a custom-built system comprising a calibrated titanium cantilever (length: 10 cm; diameter: 2 mm; spring constant: 13.27 mN/m) mounted on a vertical translation stage. Cylindrical hydrogel samples (8 mm diameter, 5 mm height) were placed beneath the cantilever. To ensure uniform force distribution, a 180-µm thick glass slide with an 8 mm diameter was positioned between the cantilever and the sample. The displacement of the cantilever (dz) was measured in real-time using a high-resolution camera equipped with a multifocus x12 objective (Navitar). The reaction force (Fr) exerted by the hydrogel was calculated using Hooke’s law: Fr=k.dz, where k is the spring constant of the cantilever. At equilibrium, this reaction force equates to the compressive force (Fc) applied to the sample. The Young’s modulus of the hydrogel was then derived by correlating the applied force (Fc) with the resultant deformation of the sample, following Hooke’s law for elastic deformation. This methodology ensured precise quantification of the mechanical properties of the hydrogels, providing reliable data to assess their suitability for replicating tissue-specific stiffness in the microphysiological system.

### 3D design and manufacture of the crypt mold

Molds for hydrogel patterning contain pillars whose dimensions correspond to the average dimension of human colonic crypts (100µm diameter and 400µm height, supplementary figure 2) [30]. These inverted patterns (pillars) allow the formation of invaginations when replicated into a scaffold. The dimensions of the crypts patterns on the 3D mold have been validated after fabrication (98 µm-diameter and 417 µm-height, Figure S1). Briefly, mold and chamber’s frame designs were made with CAO software (Fusion 360) before exported to STL file. This file was printed by stereolithography using DWS29J+ system and DS-3000 biocompatible resin from DWS Company (Italy). Regarding the parameters employed for the production, the design was sliced in 50 μm thick layers and printed with a 20 μm spot size (405 nm wave length) at 5800 mm/s speed. After printing, pieces were removed from the building table, rinsed with ethanol and then developed by an ultrasound bath during 3 minutes (P-Series Ultrasonic cleaner, Fisherbrand™). Pieces were then rinsed once again with ethanol and dried, before being post-polymerized in a UV chamber (32mW) during 30min. Additionally, molds were treated with an anti-adhesive coating by SAM (Self-Assembled Monolayer) grafting in the gas phase using SiO2 (5 min) followed by FDTS (1H,1H,2H,2H-Perfluorodecyltrichlorosilane) (5 min). This treatment forms a fluoro-silane layer between 10 and 20 nm thick, reducing adhesion and aiding in the clean release of the molded hydrogel. Molds can be used to create crypts on silanized glass slide or in the MPS.

### MPS assembly and crypts molding

The microphysiological system (MPS) comprises a stainless-steel chamber frame, fabricated by precision machining, onto which two glass coverslips—previously silanized using APTES and glutaraldehyde as described—are affixed on each side with the treated surface facing inward (Figure 2A). A thin layer of polydimethylsiloxane (PDMS; 10:1 silicone to curing agent ratio, Corning) was used as an adhesive, and the assembly was cured at 79°C for 2 hours. Metal needle tips (14G x 0.25”; Poly Dispensing Systems, Z511114) were press-fitted into the inlets and outlets of the chamber frame to serve as microfluidic connectors. These were then linked to external Tygon ND-100-80 tubing (1.78 mm outer diameter; Fisher Scientific, 15385372) for fluidic interfacing. To define the basal microfluidic channel, an internal tubing (1 mm diameter; Merck, Z609706-1PAK) was inserted through the basal port and later removed following hydrogel polymerization. Assembled chips were sterilized using a UV lamp for 1 hour prior to hydrogel molding. For crypt molding (Figure 2B), 450 µL of the prepared polyacrylamide (PA) hydrogel solution was poured into the assembled and sterilized device. The molding insert—fabricated as described earlier—was first pre-coated with 100 µL of collagen type I solution and incubated at 37°C for 30 minutes It was then carefully inserted into the chamber until it contacted the PA solution surface. After 30 minutes of gelation at room temperature, the mold was removed vertically to preserve the crypt geometry. The remaining chamber volume was filled with sterile 1X phosphate-buffered saline (PBS, pH 7.4; Sigma, D8537). The molded device (Figure 2C) was stored in 1 PBS containing 1% v/v Pen Strep (Gibco, 15140-122) + 1% v/v Amphotericin B (stock solution 250µg/ml, Gibco, 15290-026) + 0,1 % gentamicin (Sigma-Aldrich, G1397), at 4°C until further use.

### MPS preparation for culture

After 24 hours, the EnView device is closed with a PDMS lid glued with dental silicone (picodent twinsil® speed 22). We proceed to connect the tubing previously cleaned with ethanol and then PBS containing 1% v/v Pen Strep (Gibco, 15140-122) + 1% v/v Amphotericin B (stock solution 250µg/ml, Gibco, 15290-026) + 0,1 % gentamicin (Sigma-Aldrich, G1397). The chips are then filled with culture media via the microfluidic set up. A last UV bath for 30 min is done, prior to fresh media feeding into the chamber. The chips are kept at 37°C in a 5% CO2 atmosphere in the incubator 1 night before cell seeding. Cells are seeded at a density of 35 thousand cells per MPS, using a needle 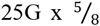 “(BD Microlance™, 240316) that goes through the PDMS lid.

### Crypts dimensions characterization

The dimensions of the crypts were determined from bright-flied confocal hydrogels images. Height was measured, in the middle of the crypt, from the bottom line to the surface of the hydrogel. Average crypt diameter was determined by measuring diameters at three positions in the crypt (bottom, middle and top). Measurements were made using ImageJ 1.53.

### Diffusion characterization

Diffusion coefficient in hard and soft polyacrylamide hydrogels was experimentally determined by perfusing from the basal channel a range of fluorescein isothiocyanate (FITC) labelled dextran solutions (weight-averaged molecular mass 3kDa (Invitrogen, D3305), 70 and 150kDa; (Sigma, 53471 and 74817 respectively). PBS alone was added into the luminal side, so that the FITC-dextran concentration difference across the hydrogel was constant thought the experiment. FITC-dextran was dissolved in PBS, and measurements were made at 0.1 mM, at 100µL/min. Diffusion through the gel was observed using confocal microscopy (Apotome 2, Zeiss) over time (image acquisition every 10 seconds), using a 5x (ON: 0.15) inverted objective. Diffusion coefficients were then calculated according to Fick’s diffusion law (Eq.1), were C* is the median concentration, C2 the final concentration, *erfc* is the error function, x is the distance of the diffusion front, D is the diffusion coefficient, t is the time and K is a constant. As the *erfc* is equal to a constant its argument must be another constant (K’) of value approximatively 1 (Eq.2). Therefore, by applying the previous equation at the initial and final time (t1 and t2, respectively) and then subtracting the square of Eq. 3.2 to Eq. 3.2, we obtain Eq. 4. Finally, we can rearrange the equation to determine the diffusion coefficient (Eq. 4.1).

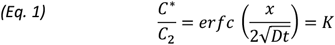

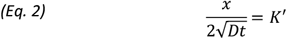

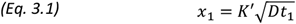

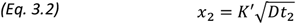

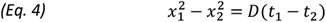

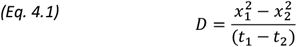

Experimentally, using ImageJ 1.53 gray profiles were measured. Gray values profiles were normalized subtracting the smallest intensity value to each one of the points (Xi –Xmin) in order to obtain values starting from 0. Then all of the values were divided by the highest intensity value to obtain values going up to 1 (Xi/Xmax). Distance from the basal channel to the diffusion front at the middle of the gray values profiles (gray value = 0.5) was then measured at the beginning and at the end of the experiment.

### Cell culture

Caco-2 cells (ATCC, htb-37) were amplified in 2D culture before seeding on the MPS. Cells were cultured in 25 cm2 culture flasks (ThermoFisher Scientific, 156367) at 37°C, 5% CO2, in DMEM 4.5 g/L glucose (ThermoFischer, 31966047) supplemented with 10% Fetal Bovine Serum (FBS, ThermoFischer, 10270), 1% non-essential amino acids (NEAA, ThermoFischer, 11140035) and 2 μg/mL penicillin/streptomycin (Sigma, P4333). Cells were passaged using 0.05% trypsin + 0.53 mM ethylenediaminetetraacetic acid (EDTA) (Sigma, T3924) every week. Cell culture experiments on the MPS were performed 2 days after hydrogels casting (at least an overnight incubation in PBS is required). Cells were seeded at 35 thousand cells per cm^2^ onto the device, 24h after sedimentation cells excess was removed by perfusion, and medium was renewed every day.

### Fluorescence staining

Cells were rinsed with PBS and fixed with 3.7% PFA (paraformaldehyde, Sigma, 252549) for 3 min. Cells were permeabilized using PBS-0.1% Triton X-100 (Sigma, T8787) for 20 min, then washed three times in PBS (PBS, Sigma, D8537). Phalloidin 568 (1:300, Invitrogen, A12380) and DAPI (1 µg/mL, Invitrogen, 62248) were added for F-actin staining and for nuclear DNA staining, respectively. Cells were incubated during 24 hours and then rinsed three times in PBS, to be further stock at 4 °C. Sample saturation, fixation, staining and washings were performed on the chip.

### Image acquisition and analysis

Brightfield images for pillar structure profilometry on the mold were obtained using a 3D digital Hirox microscope (HRX-01). During cell culture, brightfield images were captured with a Nikon Eclipse Ts2 microscope equipped with 10X or 20X objectives, a Nikon DS-Fi3 camera, and a DS-L4 control unit. Confocal imaging for hydrogel characterization was conducted on an inverted Leica SP8 confocal microscope, equipped with two PMT detectors and lasers at 488 nm, 552 nm, and 638 nm, using a 10X objective. To confirm the collagen fibrillar structure, second harmonic generation (SHG) microscopy was performed with an upright multiphoton Zeiss LSM 7 MP microscope, which included a pulsed laser adjustable from 690 to 1080 nm and a 20X water immersion objective. Confocal images of fluorescent samples were acquired using a Leica SP8 up right confocal microscope, using a 25X physio objective. Brightfield and green fluorescent time-series images for diffusion characterization were captured using an Apotome Zeiss system, using a 5X. All images were processed and analyzed using ImageJ 1.53 and IMARIS 9.5 software.

### Statistical analysis

Descriptive statistical analyses were performed and graphs were generated using the GraphPad Prism 10.1.2 Software. Data are presented as mean±SD.

## Acknowledgments

The authors thank people from the INFINITy imaging platform for assistance with biphoton imaging and people from the CBI imaging platform for assistance with confocal imaging.

## Authors contributions

Conceptualization: A.F., L.M., D.H., J.F. Hydrogel formulation: D.H., J.F., D.R-G Crypts dimension characterization: D.R-G, D.S Second Harmonic Generation on chip: D.R-G Interpenetration characterization on chip: D.R-G Diffusion assays on chip: D.R-G Cultures and immunofluorescence staining on chip: D.R-G Compression assays: S.P, D.R-G Mold pillars dimension characterization: J.F, D.H Supervision: A.F., L.M. Writing—original draft: D.H., D.R-G, A.F., L.M. Writing—review & editing: All authors contributed to the final version and editing of the manuscript Funding acquisition: A.F., L.M

## Funding

This work has benefited from several grants: Plan Cancer “System biology” 2017 and Agence Nationale de la Recherche (ANR) Molière ANR-21-CE45-0028 partly contributed to this project (granted to A.F.). The region Occitanie and Université de Toulouse III funded the salary of D.H. (granted to A.F. and L.M.). Défi-Clé BioOcc & ANR Molière ANR-21-CE45-0028 funded the salary salary D.R.G. granted to A.F.). The work was also supported by the GIS FC3R (grant agreement number R23049BS, granted to A.F.). The work was supported LAAS-CNRS micro and nanotechnologies platform, a member of the French Renatech network. It was partly supported as part of the MultiFAB project funded by FEDER European Regional Funds and French Région Occitanie (grant agreement number 16007407/MP0011594) and by the HoliFAB project funded by the European Union’s Horizon 2020 research and innovation program (grant agreement No. 760927). This work has benefited from a government grant managed by the Agence Nationale de la Recherche under the France 2030 program, reference ANR-24-EXME-0002, granted to A.F. and L.M.

### Data and materials availability

All data are available in the main text or the supplementary materials.”

## Supplementary Materials

**Supplementary figure 1.**
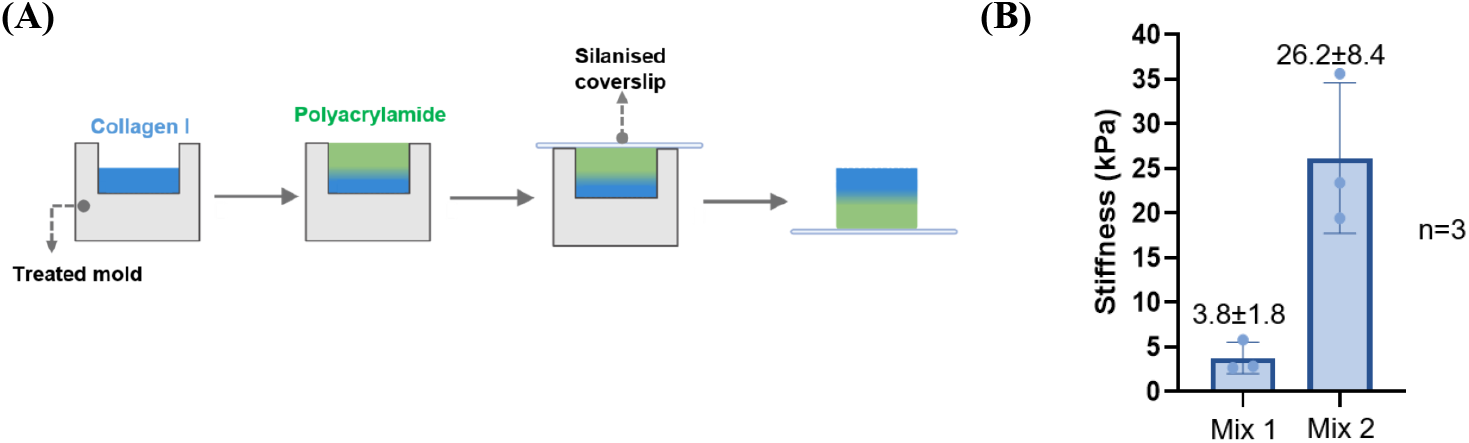
Bulk stiffness characterization of IPN hydrogels. (A) Illustration of the steps used to produce the IPN hydrogels for bulk stiffness measurements. (B) Collagen I was prepared at 5mg/ml and reticulated at 37°C. Two mixes of polyacrylamide were prepared changing acrylamide:bis -acrylamide ratio; mix 1 (1: 1.12) and mix 2 (1: 0.45) (n=3).

**Supplementary figure 2.**
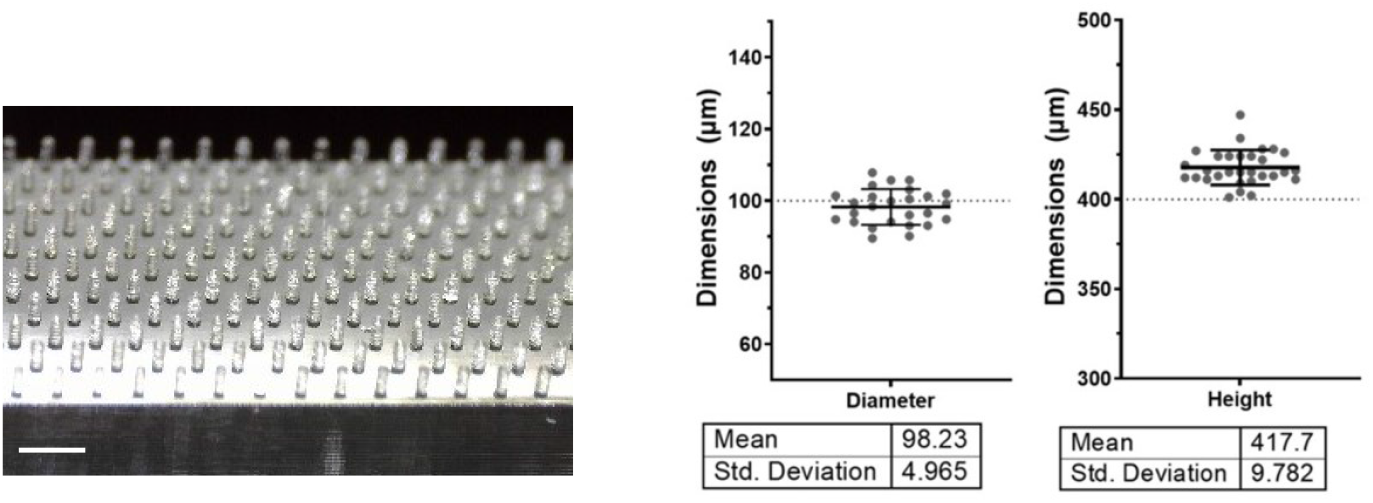
Mold characterization. After mold 3D printing, dimension of the pillars was checked to ensure if inverted crypt-like structures are as expected (100µm diameter and 400µm height). Left: bright-field representative image of the pillars on the mold (Hirox microscope, scale bar: 1mm). Right: diameter and height pillars quantification (a dotted line represents expected dimensions). Table presenting Mean ± SD values for diameter and height of the pillars (measurement done on three different molds).

**Supplementary figure 3.**
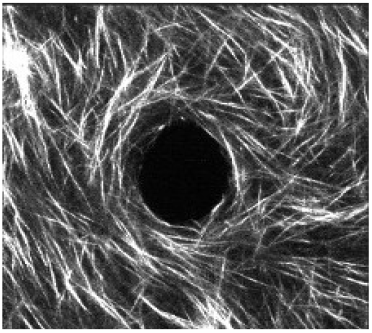
Visualization (Top view) by second harmonic imaging of the fibrillar organization of the Coll I network on the plateaus of the molded 26.2kPa IPN hydrogel. Scale bars: 100 µm

